# SARS-CoV-2 Omicron BA.1 variant infection of human colon epithelial cells

**DOI:** 10.1101/2022.04.13.487939

**Authors:** Avan Antia, David M. Alvarado, Qiru Zeng, Deanna L. Davis, Matthew A. Ciorba, Siyuan Ding

## Abstract

Omicron B.1.1.529 became the predominant SARS-CoV-2 variant in early 2022, causing a new wave of public anxiety. Compared to the ancestral strain, Omicron has 50 mutations, with over 30 mutations in the spike protein. These differences likely underlie the changes in Omicron biology noted in other studies, including an attenuation in the lung parenchyma, compared to the ancestral SARS-CoV-2 strain and other variants, as well as a preference for endosomal entry, in place of the TMPRSS2-mediated membrane fusion pathway. This raises questions on Omicron tropism and infectivity in various target organ systems, including the gastrointestinal (GI) tract. Up to 70% of COVID-19 patients report GI symptoms, including nausea, vomiting, and diarrhea. Here, we show that in the context of donor intrinsic genetic heterogeneity, the SARS-CoV-2 Omicron variant infects human colonoids similarly, if not less effectively, than the ancestral WT (WA1) strain or the Delta variant. Additionally, we note a higher ratio of viral RNA to infectious virus titer, which may suggest that Omicron is potentially less infectious in the intestine. This study lays the foundation for further work defining mechanisms mediating intestinal infection and pathogenesis by Omicron.

---

Omicron B.1.1.529 (including BA.1, BA1.1, and BA.2 subvariants) became the predominant SARS-CoV-2 variant in early 2022, causing a new wave of public anxiety. Omicron has 50 mutations compared to the ancestral strain, with over 30 mutations in the spike protein. These differences likely underlie the attenuated replication of Omicron in the lung parenchyma when compared to the ancestral SARS-CoV-2 strain and other variants^1^. Omicron may also preferentially use endosomal entry over the TMPRSS2-mediated plasma membrane fusion pathway, raising further questions on Omicron tropism and infectivity^2, 3^. It is crucial to understand the virulence and host immune responses of Omicron in various target organs, including the gastrointestinal (GI) tract. Up to 70% of COVID-19 patients experience GI symptoms, including nausea, vomiting, and diarrhea. We and others have previously shown that human small and large intestines express high levels of ACE2 and TMPRSS2/4, the host receptor and proteases, respectively, required for SARS-CoV-2 cell entry into host cells^4^. Here, we show that in the context of donor intrinsic genetic heterogeneity, the SARS-CoV-2 Omicron variant infects human colonoids similarly, if not less effectively, than the ancestral WT (WA1) strain or the Delta variant. This study lays the foundation for further work defining mechanisms mediating intestinal infection and pathogenesis by Omicron.

All study procedures and reagents were approved by the Washington University IRB (#202011003). Primary colon epithelial cells (colonoids) were derived from healthy donor biopsies and cultured as previously described^5^. Each SARS-CoV-2 isolate and passage was confirmed by RNA sequencing (**Supplemental Table S1**). Supernatant from infected transwell colonoid monolayers was titrated by focus forming assay. Fixed monolayers were stained for SARS-CoV-2 nucleocapsid (N), actin, and DAPI prior to confocal imaging. Expression levels of SARS-CoV-2 N, GAPDH, interferon lambda (IFNL3), interferon beta (IFNB), and MX1 were quantified by RT-qPCR (primers and probes in **Supplemental Table S2)**. HEK293-hACE2-TMPRSS2 cells were transfected with plasmids encoding variant spike proteins and plasmids encoding GFP and imaged at 24 hours post-transfection for syncytia formation. HEK293-hACE2 cells were transfected with plasmids encoding variant spike proteins and plasmids encoding empty vector control or V5-tagged host proteases TMPRSS2 or furin, and analyzed for spike cleavage by western blot at 24 hours post-transfection. Additional methodology is found in the **Supplementary Methods**.

Colonoids derived from four individual donors were seeded onto 2D transwell monolayers, differentiated, and infected apically with WT, Delta, or Omicron (MOI = 0.01 for 24 hours). Compared to WT and Delta, Omicron showed significantly increased replication as measured by intracellular viral RNA levels in 211A and 251A (**Fig. 1A**). Despite inter-individual differences in infectivity with each variant, a similar trend was observed in the colonoids of donor 262A (**Fig. 1A**). We conducted immunofluorescence to visualize intracellular SARS-CoV-2 N antigens in infected colonoids to confirm active replication (**Fig. 1B**). We additionally performed a focus forming assay to measure the amount of infectious SARS-CoV-2 progenies secreted into the apical colonoid supernatants, which demonstrated that Omicron produced comparable or numerically lower levels of infectious viruses than Delta and WT (**Fig. 1C**). This higher ratio of viral RNA to infectious virus titer suggests that Omicron is potentially less infectious in the intestine. Omicron also induced variable, but statistically similar, levels of type III IFN (IFNL3) expression, compared to the other SARS-CoV-2 strains (**Fig. 1D**). There was little induction of type I IFN (IFNB) and MX1, a canonical interferon-stimulated gene highly induced by type I and III IFNs, at 24 hours post-infection (**Fig. S1A**).

**Figure 1.**
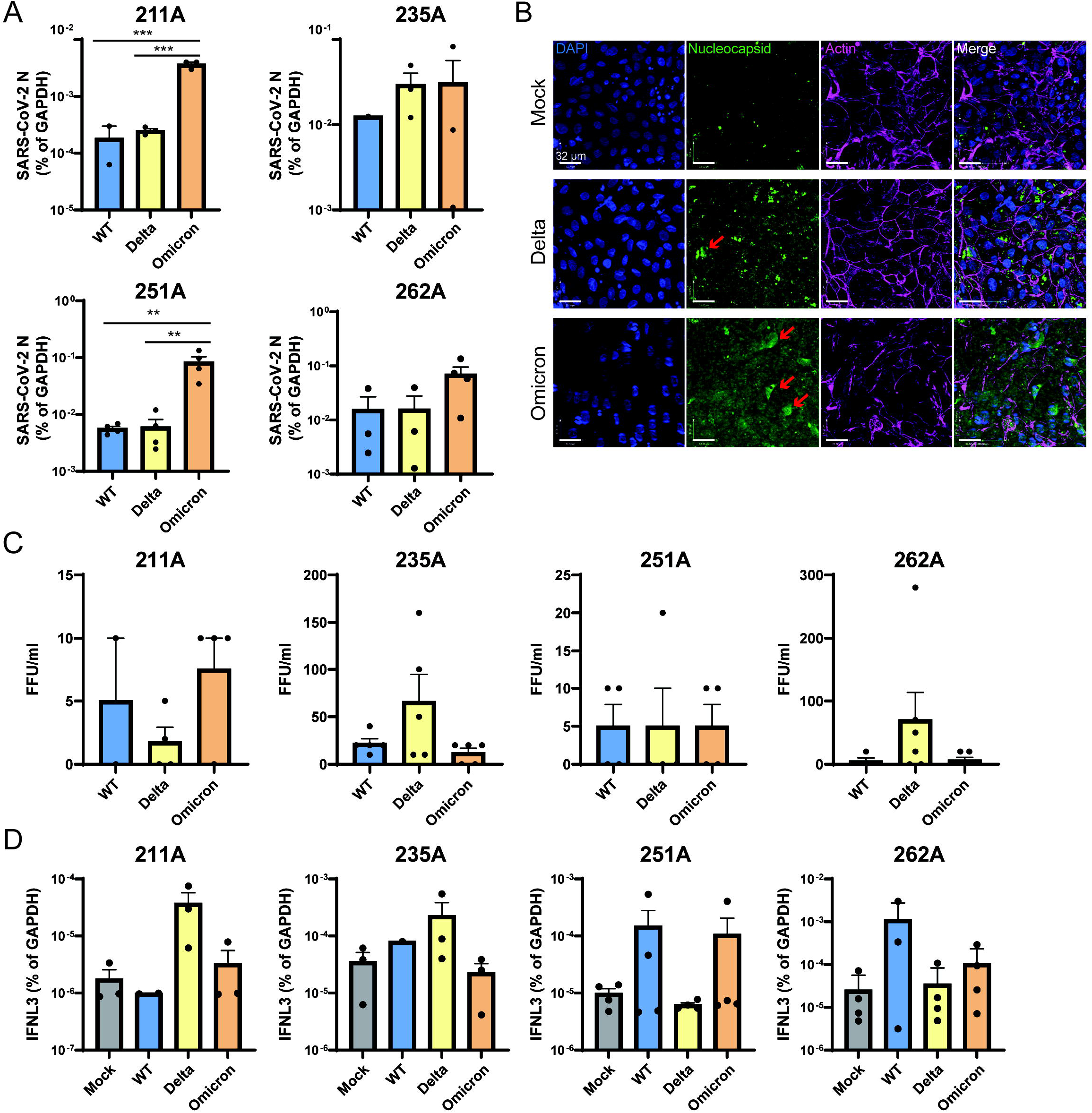
SARS-CoV-2 infection of donor derived colonoids. **(A)** Colonoid lines in 2D transwell monolayers derived from four donors were infected with indicated SARS-CoV-2 variants at an MOI of 0.01. RNA was harvested at 24 hours post infection and SARS-CoV-2 N level was quantified by RT-qPCR and normalized to that GAPDH. (Mean with SEM, One-way ANOVA with Tukey’s multiple comparisons test. ** P <0.01, *** P<0.001). (B) Colonoid 262A monolayers were infected by Delta or Omicron at an MOI of 0.01 and fixed at 24 hours post infection. Z-stacked confocal microscopic images were acquired and stained for SARS-CoV-2 N (green), actin (violet), and DAPI (blue). Merge image demonstrates intracellular viral nucelocapsid staining. Red arrows indicate intracellular viral staining. Scale bar: 32μm. (C) Quantification of infectious viral particles in the supernatants collected from the apical compartments of corresponding transwells at 24 hours post infection by a focus forming unit assay. (D) Same as **(A)** except that cellular IFNL3 mRNA level was measured instead (Mean with SEM).

To understand the molecular basis of high viral RNA and low virus titer of Omicron in human intestinal epithelial cells, we tested whether Omicron has more effective cell-to-cell spread. We ectopically expressed SARS-CoV-2 variant spike proteins in HEK293 cells stably expressing human ACE2 and TMPRSS2^6^. Omicron spike induced the formation of fewer syncytia than either WT or Delta (**Fig. S1B**), consistent with a recent study^2^. Instead, we found inefficient Omicron spike cleavage by TMPRSS2 and furin proteases (**Fig. S1C**), suggesting possible attenuation of Omicron upon viral egress, when processed into mature infectious viruses.

In this study, we found that the SARS-CoV-2 Omicron BA.1 variant effectively infects healthy donor-derived colonoids, producing high levels of intracellular viral RNA in some donors, but comparably lower levels of infectious particles. We also found that Omicron induced weak IFN response after 24 hours, possibly due to reduced recognition by cytosolic sensors or viral antagonism of IFN responses. To date, only one study has compared infectivity of SARS-CoV-2 variants in human enteroids. Using spike-pseudotyped lentiviral viruses and a luciferase-based reporter assay to quantify infection, the authors observed a 2.5- and 5-fold higher infection of colonoids with the Omicron pseudotype spike compared to Delta and D614G spikes, respectively^4^. Our study’s use of authentic SARS-CoV-2 virus and quantitation of both viral RNA and infectious particles may help explain this potential discrepancy.

GI symptoms in COVID-19 patients are strikingly frequent and have generated great interest for understanding how SARS-CoV-2 interacts with intestinal physiology. Disease states and commonly prescribed anti-inflammatory drugs can modulate intestinal ACE2 and protease expression, which potentially altered infectivity and disease severity in the initial waves^5, 7, 8^. Further, COVID-19 causes gut microbial dysbiosis and microbial diversity does not recover to pre-infection levels, even 6 months post-initial infection^9^. It is posited that such dysbiosis may be one potential contributor to long COVID-19. It is now of great interest to examine these possibilities in further studies in the context of Omicron and other SARS-CoV-2 variants. Due to the presence of viral RNA in stool and wastewater, there was concern for potential fecal-oral SARS-CoV-2 transmission^10^. Here, we show that at least in the colon, the Omicron variant does not produce more infectious particles, potentially reducing concern for GI virus shedding. In summary, our study establishes SARS-CoV-2 variant specific replication differences in human colonoids. As the pandemic evolves, there is already evidence for future variants and recombinants that can have unique features of transmission and pathology. As a potential viral reservoir, it is crucial to understand the molecular mechanisms of Omicron infection in the intestines, to better prepare for the return to normalcy.

## ACKNOWLEDGEMENTS

We extend our deepest gratitude to Naomi Sonnek of the Precision Animal Models and Organoids Core at Washington University in St. Louis, who produced all organoids and Transwells, as well as Maritza Nieves Puray Chaves, Marjorie Cornejo Pontelli, Hung Vuong, and Sebla Kutluay from Washington University in St. Louis for their help, without which this work would not have been possible. This publication was made possible in part by Grant Number UL1 RR024992 from the NIH-National Center for Research Resources (NCRR).

