## Supplemental Material for "SARS-CoV-2 Omicron BA.1 variant infection of human colon epithelial cells"

### SUPPLEMENTARY METHODS:

*Colonoid handling, infection, harvesting*: Colonoids were seeded into transwells and maintained for 7 days in 50% L-WRN conditioned medium containing 10 µM Y-27632. Differentiation was performed using Dulbecco's modified Eagle medium/F12 supplemented with 20% FBS, L-glutamine, penicillin/streptomycin, and 10 µM Y-27632 for 3 days before infection. Infections were conducted apically at an MOI of 0.01 for 1 hour at 37°C, after which viral inoculum was removed, replaced with differentiation media and incubated for 24 hours. For viral RT-qPCR quantification, cell lysates were harvested in Trizol and RNA was extracted according to manufacturer's protocol.

*Viral propagation and sequencing*: All virus passages were done in Vero E6 TMPRSS2 cells as previously described[1]. Viral stock was harvested in Trizol and RNA was extracted according to manufacturer's protocol. SARS-CoV-2 sequences were enriched using the ARTIC v4.1 primer set for SARS-CoV-2 viral enrichment and sequenced on the Illumina NovaSeq platform. Output sequences were trimmed and aligned with the SARS-CoV-2 reference (NC_045512.2) by The Genome Technology Access Center at Washington University in St. Louis.

*Immunofluorescence*: Transwells were fixed in 4% paraformaldehyde (PFA) for 20 minutes at room temperature and stained with anti-SARS-CoV-2 N antibody (40588-T62; Sino Biological), phalloidin (Alexa Fluor 647), and DAPI for immunofluorescence confocal imaging.

*Focus Forming Assay*: Vero E6-TMPRSS2 cells were seeded at $2.5 \times 10^4$ cells/well in 96-well plates and grown overnight in Dulbecco's Modified Eagle Medium supplemented to contain 10% heat-inactivated fetal bovine serum, 10 mM HEPES, and 100 U/mL penicillin/100 and U/mL streptomycin to reach confluency. Cells were transferred to a biosafety level 3 (BSL-3) facility for infection with viral supernatant collected from the apical compartment of infected transwells. Cells were incubated with 100 µl of viral supernatant at 37°C for 1 hour, after which viral supernatant was removed, replaced with 100 µl of a prewarmed overlay mixture of 2X MEM + 4% FBS with 2% methylcellulose in a 1:1 ratio, and incubated for 30 hours at 37°C[1]. After overlay removal, cells were washed 6 times with PBS and fixed with 4% PFA for 20 min, before removal of PFA and replacement with PBS. Plates were removed from BSL-3 facility and permeabilized with PBS + 0.1% Triton-X100 for 10 minutes at room temperature, washed twice with PBS + 0.1% Tween-20 (PBST), and blocked with PBST with 1% BSA and 10% FBS for 1 hour at room temperature. Upon removal of blocking buffer, cells were incubated overnight at 4°C with a primary anti-SARS-CoV-2 N (40588-T62; Sino Biological) antibody (1:1000 dilution in PBST + 1% BSA). Cells were later washed in PBST, incubated for 45 minutes at room temperature with secondary Goat anti-Rabbit IgG (Heavy chain) Superclonal Recombinant Secondary Antibody, HRP (Thermo Fisher Scientific A27036) (1:1000 dilution in PBST + 1% BSA), washed with PBST, and developed with AEC substrate kit, Peroxidase (HRP) and methods (Vector Laboratories SK-4200). Foci were quantified under an ECHO Revolve microscope.

*Syncytia assay*: HEK293-hACE2-TMPRSS2 cells were transfected using Lipofectamine 3000 reagent and manufacturer's methods (Thermo Fisher L3000015) with plasmids encoding variant spike proteins (WT pTwist-SARS-CoV-2 Δ18, plasmid #164436; pTwist-SARS-CoV-2 Δ18 B.1.617.2v1, plasmid #179905; pTwist-

SARS-CoV-2 Δ18 B.1.1.529, plasmid #179907) and plasmids encoding EGFP-N1 and imaged at 24 hours post-transfection for syncytia formation.

*Spike cleavage assay*: Plasmids encoding variant spike proteins (WT pTwist-SARS-CoV-2 Δ18, plasmid #164436; pTwist-SARS-CoV-2 Δ18 B.1.617.2v1, plasmid #179905; pTwist-SARS-CoV-2 Δ18 B.1.1.529, plasmid #179907) and plasmids encoding either EGFP-N1 (control), pcDNA3.1/nV5-TMPRSS2[2], or pLenti6.3/V5-furin host proteases (generated in-house via Gateway cloning) were co-transfected into HEK293-hACE2 cells. Lipofectamine 3000 reagent and manufacturer's methods (Thermo Fisher L3000015) were used for all transfections. At 24 hours post-transfection, cells were washed with PBS, lysed with RIPA buffer (Thermo Fisher Scientific 89901) supplemented with Halt Protease Inhibitor Cocktail (100X) (Thermo Fisher Scientific 78429), and incubated on ice for 10 minutes. Cell lysates were then subjected to centrifugation at 13,500 RPM for 10 minutes at 4°C to remove cell debris and nucleus. Protein samples were boiled in 2X Laemmli Sample Buffer (Bio-Rad) containing 5% β-mercaptoethanol at 95°C for 5 minutes. Prepared samples were run in 4%-12% Mini-PROTEAN TGX Precast protein gels (Bio-Rad 4561085) and transferred onto nitrocellulose membranes using Bio-rad wet/tank blotting system. Membranes were blocked in 5% BSA in TBS + 0.1% Tween-20 (TBST) at room temperature before incubation with primary antibodies at 4°C overnight. Membranes were then washed three times with TBST and incubated in secondary antibodies diluted in 5% BSA in TBST at room temperature for 1 hour. Finally, membranes were washed with TBST and visualized by using Chemi-Doc imaging system (Bio-Rad). *Primary antibodies*: SARS-CoV-2 Spike S2 Rabbit pAb (Sino Biological 40590-T62), SARS-CoV-2 Spike S1 RBD Rabbit pAb (Sino Biological 40592-T62), V5-Tag Rabbit mAb (Cell Signaling Technology 13202S), and GAPDH (BioLegend 631402). *Secondary antibodies*: Goat anti-Rabbit IgG (Heavy chain), Superclonal Recombinant Secondary Antibody, HRP (Thermo Fisher Scientific A27036).

*Statistical Analysis*: All data were subjected to Shapiro-Wilk test for normality and were subjected to parametric or non-parametric analysis of variance (ANOVA) as appropriate. Statistics were performed using Graphpad Prism 9.

**Table S1: Virus sequencing results: Spike Mutations in SARS-CoV-2 variants sequenced by NovaSeq**

| Omicron B.1.1.529 | Delta B.1.617.2 | WT WA1 |
|---|---|---|
| T95I, G142D, V143del, Y144del, Y145del, N211I, L212del, G339D, S371L, S373P, S375F, K417N, N440K, G446S, S477N, T478K, E484A, Q493R, G496S, Q498R, N501Y, Y505H, T547K, D614G, H655Y, N679K, P681H, N764K, D796Y, N856K, Q954H, N969K, L981F, D1146 | T19R, G142D, E156G, F157del, R158del, L452R, T478K, D614G, P681R, F855S, D950N, C1235F | No spike mutations |

**Table S2: qPCR Primers and Probes**

| Primer name | Sequences |
|---|---|
| SARS-CoV-2 N | Forward: ATGCTGCAATCGTGCTACAA |

| | |
|---|---|
| | Reverse: GACTGCCGCCTCTGCTC<br>Probe: FAM/TCAAGGAACAACATTGCCAA/TAMRA |
| Human GAPDH | Forward: GGAGCGAGATCCCTCCAAAAT<br>Reverse: GGCTGTTGTCATACTTCTCATGG |
| IFNL3 | Forward: TAAGAGGGCCAAAGATGCCTT<br>Reverse: CTGGTCCAAGACATCCCCC |
| IFNB | Forward: ATGACCAACAAGTGTCTCCTCC<br>Reverse: GGAATCCAAGCAAGTTGTAGCTC |
| MX1 | Forward: GTGGCTGAGAACAACCTGTG<br>Reverse: GGCATCTGGTCACGATCCC |
