## Supplementary figures and images for "SARS-CoV-2 Omicron BA.1 variant infection of human colon epithelial cells"

### Supplemental Figure 1

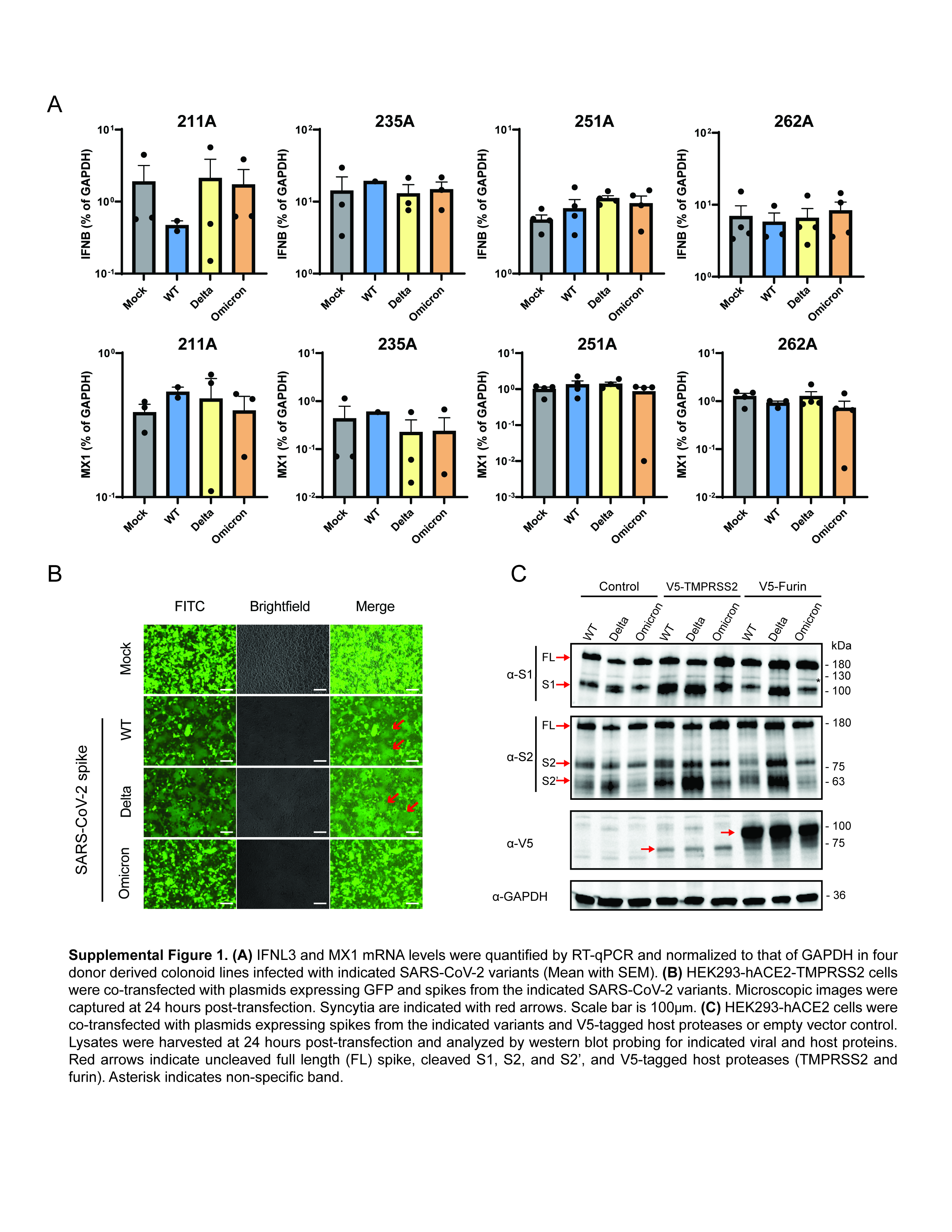
